# Essential Roles of the *SPPRA* Fructose-Phosphate Phosphohydrolase Operon in Carbohydrate Metabolism and Virulence Expression by *Streptococcus Mutans*

**DOI:** 10.1101/342170

**Authors:** Lin Zeng, Robert A. Burne

## Abstract

The dental caries pathogen *Streptococcus mutans* can ferment a variety of sugars to produce organic acids. Exposure of *S. mutans* to certain non-metabolizable carbohydrates such as xylitol impairs growth and can cause cell death. Recently, the presence of a sugar-phosphate stress in *S. mutans* was demonstrated using a mutant lacking 1-phosphofructokinase (FruK) that accumulates fructose-1-phosphate (F-1-P). Here we studied an operon in *S. mutans, sppRA*, which was highly expressed in the *fruK* mutant. Biochemical characterization of a recombinant SppA protein indicated that it possessed hexose-phosphate phosphohydrolase activity, with preferences for F-1-P and, to a lesser degree, fructose-6-phosphate (F-6-P). SppA activity was stimulated by Mg^2+^ and Mn^2+^, but inhibited by NaF. SppR, a DeoR-family regulator, repressed the expression of the *sppRA* operon to minimum levels in the absence of the fructose-derived metabolites, F-1-P and likely also F-6-P. Accumulation of F-1-P, as a result of growth on fructose, not only induced *sppA* expression, it significantly altered biofilm maturation through increased cell lysis and enhanced extracellular DNA release. Constitutive expression of *sppA*, via a plasmid or by deleting *sppR*, greatly alleviated fructose-induced stress in a *fruK* mutant, enhanced resistance to xylitol, and reversed effects of fructose on biofilm formation. Finally, by identifying three additional putative phosphatases that are capable of promoting sugar-phosphate tolerance, we show that *S. mutans* is capable of mounting a sugar-phosphate stress response by modulating the levels of certain glycolytic intermediates, functions that are interconnected with the ability of the organism to manifest key virulence behaviors.

**Importance:** *Streptococcus mutans* is a major etiologic agent for dental caries, primarily due to its ability to form biofilms on tooth surface and to convert carbohydrates into organic acids. We have discovered a two-gene operon in *S. mutans* that regulates fructose metabolism by controlling the levels of fructose-1-phosphate, a potential signaling compound that affects bacterial behaviors. With fructose becoming increasingly common and abundant in the human diet, we reveal the ways fructose may alter bacterial development, stress tolerance, and microbial ecology in the oral cavity to promote oral diseases.

## Introduction

Most bacteria use the sugar:phosphotransferase system (PTS) to internalize mono- and di-saccharides, a process that uses phosphoenolpyruvate (PEP) to energize the transport and concomitant phosphorylation of the carbohydrates (1). While certain components of the PTS are vital to the growth and persistence of many bacteria, the accumulation in cells of phosphorylated carbohydrates can lead to sugar-phosphate stress, a condition that often manifests in growth inhibition or cell death (2-4). A number of mechanisms have been proposed to contribute to sugar-phosphate stress: (i) futile cycling, in which the cells expend energy to internalize and phosphorylate the sugar, but the sugar is then dephosphorylated and expelled (5); (ii) reduction of essential metabolic intermediates arising from an inability to internalize other carbohydrates (6); (iii) creation of toxic metabolic byproducts, such as methylglyoxal (7, 8), or (iv) deleterious effects of specific sugar phosphates that act as allosteric modifiers of enzymes or regulatory proteins, causing dysregulation of metabolism (9). The majority of research on sugar-phosphate stress has been conducted with Gram-negative, model organisms, with far fewer studies carried out with known pathogens or Gram-positive bacteria (10).

*Streptococcus mutans*, a primary dental caries pathogen, is a Gram-positive, lactic acid-producing bacterium that depends on, and is particular adept at, transporting and fermenting carbohydrates for energy production and growth. The most studied sugar-phosphate stress in *S. mutans* is that induced by xylitol, a non-metabolizable pentitol that is internalized as xylitol-5-phosphate (X-5-P) by a fructose-specific PTS, but is subsequently dephosphorylated and released from the cells (11-13). Although the exact nature of the process remains to be clarified, including the enzyme(s) responsible for removing the phosphate group from X-5-P, it is generally believed that both the futile cycling of xylitol and the presence of X-5-P in cells are detrimental. While futile cycling wastes energy, it has been proposed that X-5-P is an inhibitor of phosphofructokinase that causes a block in glycolysis (9). The observed effects of xylitol intake by *S. mutans* include reductions in high-energy metabolites, such as fructose-1,6-bisphosphate (F-1,6-bP) and glucose-6-phosphate (G-6-P), as well as impaired glucose-PTS activity, decreased rates of acid production (14), and inhibition of the formation of extracellular insoluble polysaccharides from sucrose (15, 16). Given its ability to negatively affect a caries pathogen, xylitol is widely used as a non- or anti-cariogenic additive in products that include chewing gum and candies, despite the fact that spontaneous emergence of xylitol-resistant mutants can occur in habitual users of xylitol-containing products (17, 18). It has been shown that the fructose-PTS permease encoded by the *fruI* gene is responsible for transporting the majority of xylitol, and a *fruI-*deficient mutant lost sensitivity to presence of xylitol (18). Another example of sugar-phosphate-related stress in *S. mutans* is one induced by 2-deoxyglucose (2-DG), an analog of glucose that can be internalized and phosphorylated by the PTS, but cannot be metabolized. Uptake of 2-DG by *S. mutans* reduces the transport of carbohydrates via the PTS (19) and the multiple sugar metabolism (*msm*) system, an ATP-binding cassette (ABC) transporter for melibiose, raffinose, and certain other carbohydrates that are found in the human diet (20).

Fructose is transported into *S. mutans* via multiple PTS permeases and further metabolism requires phosphofructokinase enzymes, including a 1-phosphofructokinase and 6-phosphofructokinase that convert fructose-1-phosphate (F-1-P) and fructose-6-phosphate (F-6-P), respectively, into F-1,6-bP that can be further catabolized via glycolysis (3). We recently engineered a point mutant in *S. mutans* strain UA159 (*fruK*M28stop, strain FruK-13) that prevents the translation of an intact 1-phosphofructokinase (FruK) enzyme (3). Due to the accumulation of F-1-P in FruK-13, a severe growth phenotype is evident when fructose is present. FruK-13 also displayed significant growth defects when cultured in media containing glucose, sucrose, mannose, galactose, lactose or certain amino sugars, demonstrated a broad influence of F-1-P on carbohydrate metabolism and growth of *S. mutans* (3). RNA deep sequencing (RNA-Seq) was used to define changes in the transcriptome arising from loss of the FruK enzyme. The expression of 394 genes, including genes encoding ribosomal proteins, proteins required for biosynthesis of nucleotides and amino acids, and components of the PTS, were altered in FruK-13 compared to the parental strain UA159 (3). Among the most up-regulated genes in the *fruK* mutant were two open reading frames (ORF) in an apparent operonic arrangement, SMU.507 and SMU.508; designated *<u>s</u>*ugar-*<u>p</u>*hosphate *<u>p</u>*hosphatase *<u>R</u>*egulator *sppR* and the phosphatase *sppA*, respectively. SppR is a DeoR-type transcription regulator and SppA is a predicted hydrolase of the HAD (haloacid dehalogenase) family IIB that includes enzymes such as trehalose-6-phosphatase, plant and cyanobacterial sucrose-phosphatases, and a large subfamily of bacterial Cof-like hydrolases; Cof being an *E. coli* phosphatase (21). While both *sppR* and *sppA* are well conserved in *S. mutans* strains, homologous systems comprised of both genes in tandem have only been identified in *Streptococcus gordonii* and *Streptococcus sanguinis* (http://www.microbesonline.org). In this communication, we investigated the regulation and function of the *sppRA* operon and explored the contribution of its gene products to fitness and cariogenic potential of *S. mutans*, particularly in the context of regulation of F-1-P levels.

## Results

### Overexpression of *sppA* enhances growth of FruK-13 in the presence of fructose

In a previous study with a mutant lacking a 1-phosphofructokinase (*fruK*), FruK-13, an operon encoding a DeoR-like regulator (SMU.507, *sppR*) and a predicted hydrolase of the HAD superfamily (SMU.508, *sppA*) was significantly up-regulated (3). As FruK-13 displays the most severe growth defect on fructose, likely due to accumulation of F-1-P, it was posited that certain enzymes encoded by *S. mutans* could eliminate these potentially toxic intermediates. The putative sugar-phosphate phosphatase encoded by *sppA* was cloned into an expression vector pIB184 behind a constitutive promoter and the resultant plasmid, pIB508, was introduced into FruK-13. After confirming enhanced expression of *sppA* by plasmid pIB508 (Fig. 1A), these resultant strains were compared in their growth phenotypes in a fructose-based medium. Compared to the vector-only control strain FruK-13/pIB184, which grew very slowly on a TV-fructose medium, FruK-13/pIB508 grew at a significantly improved rate (Fig. 1B), providing a strong support for the hypothesis that SppA could function as a sugar-phosphate phosphatase that decreased the levels of F-1-P in the *fruK* genetic background.

**FIGURE 1.**
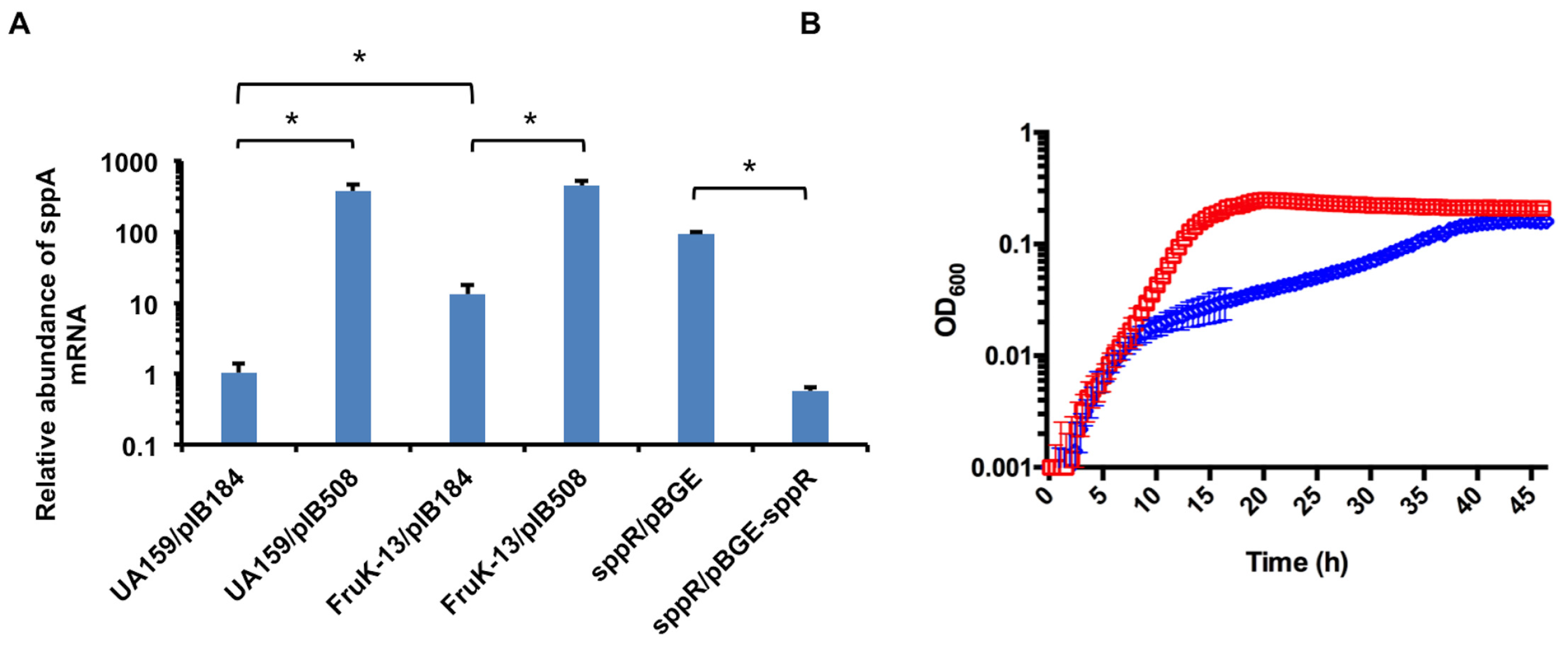
Overexpression of *sppA* enhances growth of a *fruK* mutant on fructose. (A) Relative *sppA* mRNA levels measured in UA159/pIB184, UA159/pIB508, FruK-13/pIB184, FruK-13/pIB508, *sppR*/pBGE and *sppR*/pBGE-*sppR*. Cultures were prepared using BHI medium and harvested at the exponential phase. Quantitative reverse transcription PCR (qRT-PCR) was carried out using the total RNA samples, and the relative abundance of *sppA* was determined using the ∆∆Cq method. The error bars denote the standard deviations. Student’s *t*-test was performed to determine the statistical significance of the data, with asterisks denoting *P* <0.05. (B) FruK-13/pIB508 (red squares) and FruK-13/pIB184 (blue diamonds) were cultivated in BHI and then diluted into TV medium containing 10 mM fructose. The cultures were overlaid with mineral oil and OD_600_ was measured every 30 min in a Bioscreen C. The results represent the average of at least three biological replicates.

### SppA has hexose-phosphate phosphohydrolase activity

Multiple computer algorithms predicted that SppA is a hydrolase of the HAD superfamily (e.g. BLAST, CLUSTAL). To study the function of SppA *in vitro*, the coding sequence of *sppA* was cloned into pQE30 and expression was induced in *E. coli* to produce an N-terminally His-tagged recombinant protein, designated His-SppA. His-SppA has 274 amino acids, excluding the His-tag, and migrated as a single 31-kDa band in SDS-polyacrylamide gel electrophoresis (SDS-PAGE, Fig. S1). The protein was relatively stable in a variety of buffers and pH values, although addition of 0.5 M NaCl and at least 10% of glycerol was required to retain activity during longer-term storage (>1 month) at 4°C. To test the kinetics of SppA in removing the phosphate group from presumptive substrates, a set of phosphate-free buffers were tested to carry out these experiments (see Materials and Methods for detail), and the phosphate released in each reaction was monitored using a Malachite green reagent kit.

### Substrate specificity

Using 2.5 mM *p*-nitrophenylphosphate (*p*NPP) as a general colorimetric substrate for phosphatases, phosphatase activity by His-SppA was detected at 37°C in 50 mM HEPES buffer at pH 7.5. SppA activity on *p*NPP could be stimulated by the addition of millimolar concentrations of MgCl_2_ (data not shown). We then conducted a number of preliminary experiments to gain insights into optimal conditions for activity of His-SppA on a number of sugar phosphates. Then, as determined by an end-point assay performed under optimized conditions (Fig. 2A), the preferred substrate for His-SppA was found to be F-1-P, followed by F-6-P. Very little activity was observed when F-1,6-bP, G-6-P, or GlcN-6-P were used as substrates, and SppA had no activity on GlcNAc-6-P under the conditions tested.

**FIGURE 2.**
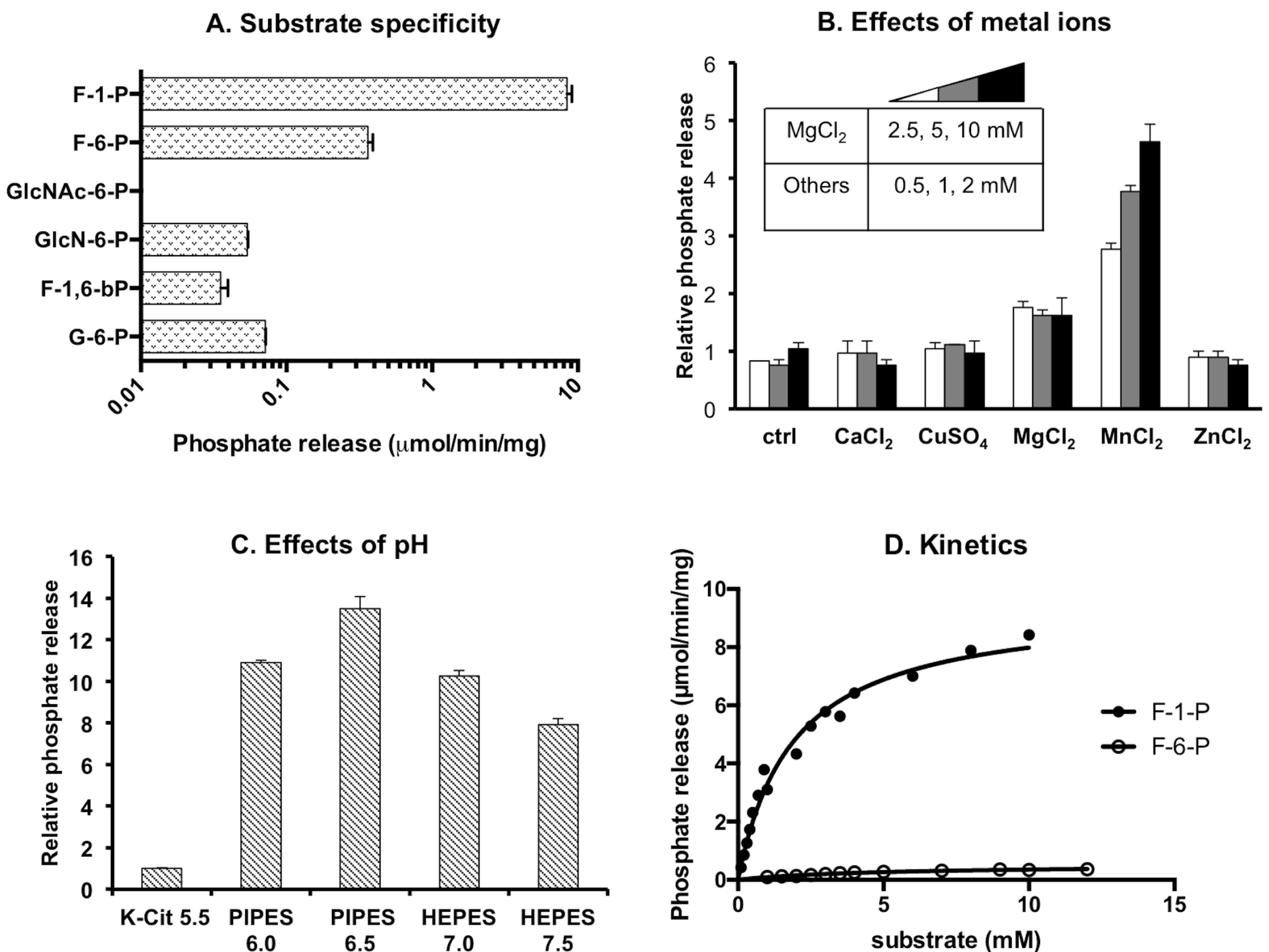
*In vitro* characterization of SppA as a hexose-phosphate: phosphohydrolase. The majority of the reactions (except panels B&C) were performed at 37°C in a basic buffer composed of 50 mM PIPES (pH 6.5), 5 mM MgCl_2_, and 1 mM MnCl_2_. Release of phosphate was measured using a Malachite Green Reagent (MGR), and the results were divided by time and amounts of enzyme (A, D), or normalized for relative activities (B, C). (A) 10 mM of the indicated sugar-phosphates were incubated with His-SppA (0.08 μM for F-1-P, 0.67 μM for the rest), with release of phosphate being monitored over a period of 10 min. (B) 0.1 mM of F-1-P was incubated for 5 min with 0.33 μM of His-SppA and the specified amounts of metal ions. (C) 0.67 mM F-1-P was incubated with 0.04 μM of His-SppA for 130 sec at the indicated pH values (See Methods for details). (D) Increasing concentrations of F-1-P and F-6-P were each mixed with His-SppA and phosphate release was monitored over defined periods of time (See Methods for details).

### Effects of metal ions

Many phosphatases are affected by the presence of metal ions (22), thus the activity of His-SppA was studied by adding to the reactions millimolar levels of CaCl_2_, CuSO_4_, MgCl_2_, MnCl_2_, and ZnCl_2_. CaCl_2_, CuSO_4_, and ZnCl_2_ at these levels (0.5 to 2 mM) had no detectable impact to the activity of His-SppA on F-1-P as a substrate, but the presence of MgCl_2_ and MnCl_2_ significantly enhanced the release of phosphate by the purified, recombinant enzyme (Fig. 2B). By testing both Mg^2+^ and Mn^2+^ together, it was determined that optimal enzymatic activities could be elicited when MgCl_2_ was present at 5 mM, along with 1 mM MnCl_2_.

### pH

The optimal pH for the activities of His-SppA was determined to be about 6.5, and the rate of phosphate release significantly decreased when the pH values of the reaction deviated from this point, particularly at values of 6.0 and below (Fig. 2C). Since *S. mutans* maintains ∆pH values of 0.5 to 1.0 units above the external environment (23), SppA is probably most active in cells growing in a moderately acidic environment, such as what would occur when *S. mutans* is actively metabolizing carbohydrate(s).

### Fluoride

Sodium fluoride (NaF) is an inhibitor of many phosphatases. The presence of 10 mM NaF significantly reduced (by >50%) the rate of phosphate release from F-1-P by His-SppA (Fig. S2 in the supplemental material).

### Reaction kinetics on F-1-P and F-6-P

Finally, using the optimal pH and metal ion conditions described above, *V_max_* and *K_m_* of His-SppA were measured using F-1-P or F-6-P. The purified enzyme was incubated with increasing concentrations of each substrate (see Materials and Methods for details) at 37°C, and release of phosphate over time was monitored by periodic sampling of the reactions. The release rate of phosphate for each reaction was then plotted against the substrate levels to estimate rate and substrate affinity (Fig. 2D). For F-1-P, the *V_max_* was 9.50 ± 0.38 μmol/(min x mg protein), and the *K_m_* was 1.90 ± 0.21 mM. For F-6-P, the *V_max_* was 0.52 ± 0.03 μmol/(min x mg protein) and the *K_m_* was 4.83 ± 0.51 mM. Thus, under these conditions, F-1-P is a preferred substrate for SppA, whereas F-6-P can be dephosphorylated by SppA, albeit less efficiently. These data are consistent with idea that *S. mutans* may use SppA to limit accumulation of F-1-P, which can be further phosphorylated (by FruK) into F-1,6-bP. However, the data may also reflect a preference of the organism to internalize then further metabolize F-6-P. These parameters, including *V_max_* and *K_m_*, are also in line with what has been reported for other members of the HAD family phosphatases in bacteria (5, 21, 24).

### SppR is a negative regulator of *sppA* that responds to fructose-phosphate signals

SppR is predicted to be a member of the DeoR family, a group of transcription regulators that often respond allosterically to metabolic intermediates including phosphorylated carbohydrates. To begin understanding the mode of regulation by SppR, a deletion mutant of *sppR* was constructed and the expression of *sppA* was measured using quantitative RT-PCR (qRT-PCR). The results showed a drastic enhancement in mRNA levels of *sppA* due to the loss of SppR, when both the wild type and the mutant were growing in the rich medium BHI (Fig. 1A and data not shown). To ensure that increased expression of *sppA* was not a result of secondary mutations, plasmid pBGE-*sppR* was created (25) and introduced into the *sppR* mutant, allowing for stable integration of the *sppR* gene in the chromosome in the *gtfA* locus. Compared to cells transformed with the empty vector, complementation of the *sppR* mutant with *sppR* in single copy resulted in expression levels of *sppA* that were comparable to those measured in the wild-type background (Fig.1A).

Since higher expression of both *sppR* and *sppA* was noted using RNA-Seq of the *fruK* mutant (FruK-13), which should have elevated levels of F-1-P, it was hypothesized that SppR negatively regulates the *spp* operon, with SppR being allosterically regulated by fructose phosphates, including F-1-P. To test this hypothesis, the promoter activity of the operon (P*sppR*) was examined by utilizing a promoter-reporter gene fusion P*sppR::cat* that was integrated in single copy at a distant site on the chromosome. Expression of the *cat* fusion was measured in cells growing on various carbohydrates. A UA159 derivative carrying the P*sppR::cat* fusion was cultivated in TV medium supplemented with 10 mM glucose to mid-exponential phase, then pulsed for 1 h with 25 mM or 50 mM fructose to increase the levels of fructose-derived metabolites. In addition, 25 mM or 50 mM of glucose, xylitol, or sucrose were also used to treat these cells. The CAT activities measured (Fig. 3A) on cells pulsed with fructose showed significant, concentration-dependent increases compared to cells pulsed with glucose. A slight reduction in CAT activity was noted in cells pulsed with 50 mM xylitol, and sucrose had little to no impact on the expression of the *spp* promoter fusion. When compared to glucose, treatment of cells with fructose should result in accumulation of F-1-P, adding support to the idea that F-1-P is an allosteric effector capable of relieving the repression by SppR. On the other hand, xylitol treatment could potentially reduce the capacity of the PTS and carbon flux, reducing the levels of glycolytic metabolites, including F-1-P and F-6-P. The lack of response from sucrose treatment could be due to the preference by the bacterium to utilize the glucose moiety, as our recent study has indicated that free fructose is expelled by *S. mutans* after sucrose is transported and phosphorylated on the glucose moiety by ScrA, and the resultant sucrose-6-phosphate (S-6-P) is cleaved by ScrB (26).

**FIGURE 3.**
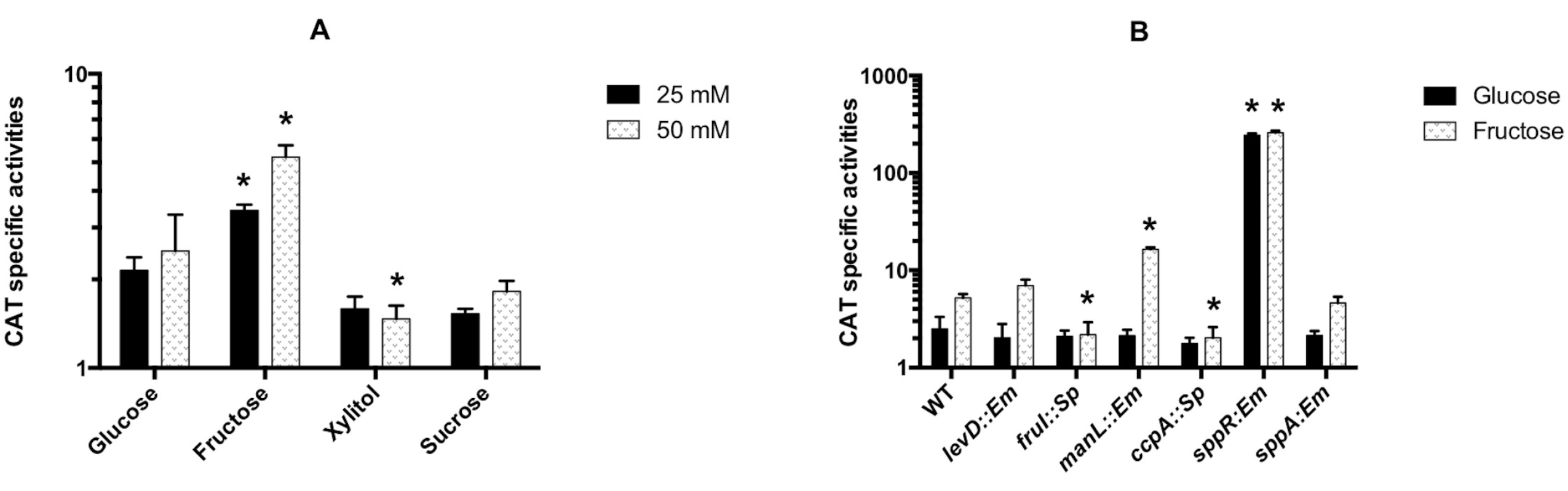
Transcription of the *spp* operon is regulated by SppR, fructose derivatives and the PTS. The P*sppR::cat* fusion was integrated into the chromosome of the wild type (A) or various mutants (B) to monitor the expression of the operon. All strains were first cultivated to mid-exponential phase (OD_600_ = 0.3 to 0.4) in a TV medium with 10 mM glucose, then (A) 25 or 50 mM of the specified carbohydrates, or (B) 50 mM of glucose or fructose was added, the cultures were incubated for 1 h, and cells were harvested for CAT assays. Error bars denote standard deviations. Asterisks indicate statistically significant difference (by Student’s *t*-test, *P* <0.05) in comparison (A) with glucose at the same concentrations, or (B) with the wild type (WT) treated with the same carbohydrates.

We have previously shown that fructose internalization via FruI likely results in creation of F-1-P, whereas internalization of fructose by EII^Lev^ (*levDEFG*) yields F-6-P (3). To test if transport of substrates via different PTS permeases could affect the regulation of expression of the *spp* operon, the P*sppR::cat* fusion was placed in the background of mutants deficient in the fructose-PTS permeases, *levD* or *fruI*, or in a mutant lacking the primary glucose-PTS permease (*manL*) (27). A mutant defective in production of catabolite control protein A (CcpA), a major regulator of genes involved in carbon flow in *S. mutans*, was also included as CcpA can regulate the expression of certain PTS permease genes (28). The wild-type control and the mutant strains were grown in TV-glucose to mid-exponential phase, then pulsed with 50 mM fructose to induce the P*sppR::cat* fusion. In support of our hypothesis that F-1-P is required for inducing expression of *spp*, loss of *fruI* abolished fructose-dependent induction of CAT activity, whereas the *levD* mutant behaved like the wild type (Fig. 3B). Interestingly, a deletion of the glucose-PTS gene *manL* resulted in significantly increased expression of the P*sppR::cat* fusion, consistent with our recent study that showed interconnected regulation of fructose and glucose PTS permeases in *S. mutans* (3). The *ccpA* mutant showed no difference in response to fructose.

To gauge the overall impact of the regulation by SppR on the *sppRA* operon and the potential that depletion of F-1-P associated with *sppA* induction could function in feedback regulation of the operon, we assayed the expression of P*sppR::cat* fusion in strains deficient in *sppR* and *sppA*, respectively. The results (Fig. 3B) showed nearly a 2-log increase in CAT activities in the *sppR* mutant regardless of the sugars tested, and no change in the *sppA* mutant. Together, these results suggest that SppR exerts a strong, negative regulation over the *spp* operon, resulting in low levels of expression of *sppA* even when fructose is present, at least in an otherwise wild-type genetic background.

### Measurements of fructose metabolites

To demonstrate that the activities of the *spp* gene products can regulate the levels of intracellular fructose-derived metabolites, bacterial cells were grown on TV-glucose to mid-exponential phase, an additional 50 mM glucose or fructose was added, the cultures were incubated for 30 min, then cells were harvested. Enzymatic assays were performed on the deproteinized cell lysates to measure F-1-P levels. The results (Table 2) showed that treatment of the wild-type strain UA159 by fructose increased F-1-P levels by ~40 fold. In comparison to the wild type, the *sppR* mutant had 3-fold lower levels of F-1-P when exposed to fructose, consistent with the fact that *sppA* is overexpressed due to the loss of its negative regulator, SppR. Under the same condition, the *sppA* mutant had similar levels of F-1-P to that of UA159. Furthermore, the *fruK* deficient strain FruK-13 produced the highest amounts of F-1-P when exposed to glucose, providing direct evidence that accumulation of F-1-P is the cause of the growth defect of FruK-13 on various carbohydrates (3). However, F-1-P levels in FruK-13 exposed to fructose appeared slightly lower than the wild type, although not statistically significant; an effect likely caused by the overexpression of *sppA*. Further, in agreement with the finding that loss of *fruI* (but not *levD*) abolished induction by fructose of the P*sppR::cat* fusion (Fig. 3B), here only the *fruI* mutant showed reduced F-1-P levels compared to UA159, when treated with fructose. When F-6-P was measured using these cell lysates, similar levels of F-6-P were detected in all four strains tested (Table 2), when each strain was exposed to either glucose or fructose.

### SppR binds to the *spp* promoter, F-6-P and F-1-P

To demonstrate the direct interaction of SppR with its own promoter region, a recombinant MBP (maltose-binding protein) -SppR was engineered and overexpressed in *E. coli*. The purified MBP-SppR was cleaved with a protease to release SppR (Fig. S1B), which was used in an electrophoretic mobility shift assay (EMSA) that included a biotin-labeled DNA fragment (*spp* probe) containing the 250-bp intergenic region immediately upstream of the *sppR* coding sequence. As shown in Fig. 4, the presence of as little as 62.5 nM SppR resulted in significant shifting of the *spp* probe. Importantly, when the reaction was repeated with the addition of 5 or 10 mM F-1-P, SppR showed significantly lower DNA-binding activities, as more probe was left unshifted (Fig. 4). Interestingly, F-6-P also showed a comparable capacity to reduce the DNA-binding activity of SppR. In comparison, F-1,6-bP had little effect on the interaction of SppR with the probe. As a negative control, a purified MBP protein was used in EMSA with the *spp* probe and no shifting of the probe was observed (data not shown).

**FIGURE 4.**
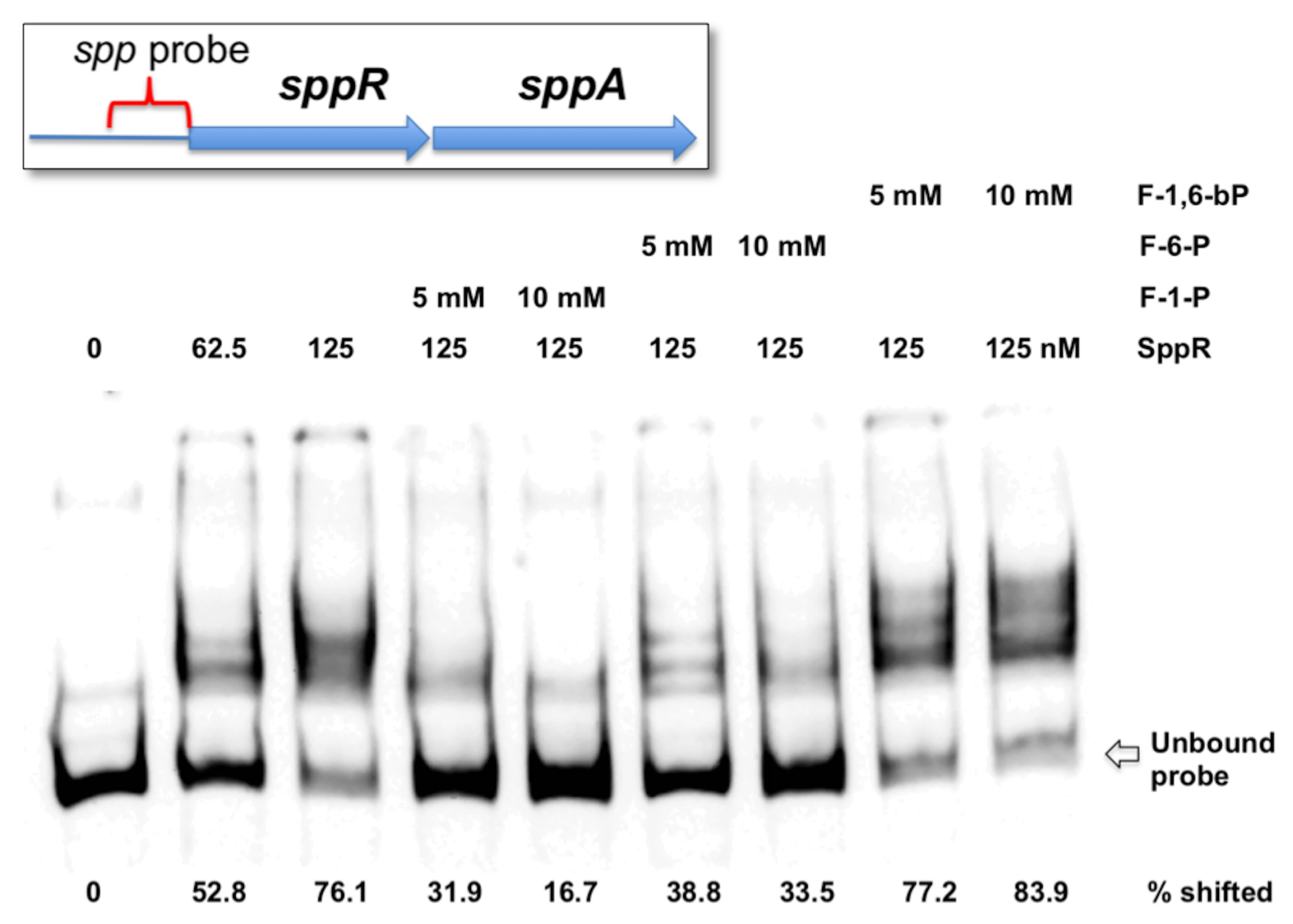
*In vitro* interactions of SppR with the *spp* promoter region and fructose phosphates. EMSA was performed using SppR and a biotin-labeled probe containing the promoter region of *sppR.* SppR (0 to 125 nM) was incubated first with various sugar phosphates for 10 min, then with 0.2 nM of DNA probe for 20 min, before the mixture was resolved on a non-denaturing polyacrylamide gel.

As an alternative to EMSA, we also performed a thermal shift assay in which increasing temperature induces unfolding of SppR, which is monitored using a protein dye that fluoresces when bound to unfolded polypeptides (29). The melting temperature (Tm) of proteins usually increases in the presence of compounds that are bound by the protein. We added to the melting reaction various amounts of glucose, G-6-P, fructose, F-1-P, F-6-P, F-1,6-bP, GlcN, GlcN-6-P, GlcNAc and GlcNAc-6-P. While most sugar and sugar phosphates had little to no impact to the unfolding of SppR, addition of F-6-P increased the Tm in a dose-responsive manner, indicating a specific interaction between SppR and F-6-P (Fig. S3 and data not shown). Meantime, addition of F-1-P did not significantly change the Tm of SppR. Considering the results of our genetic analysis, we posit that the interactions between SppR and fructose-derived metabolites modulate the repression of *spp* expression. It is intriguing that F-1-P showed little affinity for SppR in the thermal shift assay, as it was quite effective at inhibiting the binding of SppR to the *spp* promoter region. It is possible that efficient interaction of SppR to F-1-P requires the presence of its cognate DNA molecules or other factors that were present only in the EMSA reactions. In the future, we plan to utilize a spectrum of binding assays to further evaluate the possible allosteric effectors that modulate the DNA-binding affinity of SppR *in vivo.*

### Overexpression of SppA enhances resistance to xylitol

Having demonstrated that enhanced expression of SppA from pIB508 abolished the sugar-phosphate stress in strain FruK-13, we tested if overexpression of SppA could also alleviate stress caused by the presence of xylitol. Strains UA159/pIB184 and UA159/pIB508 were created and their growth was monitored in TV containing 0.5% glucose or fructose, with or without 1% xylitol. Without addition of xylitol, both strains grew similarly well in TV glucose or fructose (data not shown). When 1% xylitol was added to the TV-glucose medium, UA159/pIB184 displayed a reduced growth rate compared to when the strain was grown in TV with glucose alone. Interestingly, UA159/pIB508 showed significantly faster growth than UA159/pIB184 in the presence of both glucose and xylitol, indicating enhanced resistance to xylitol as a result of increased expression of SppA (Fig. 5A; see Fig. 1A for *sppA* mRNA levels). In contrast, addition of 1% xylitol to TV-fructose had no discernable effect on the growth phenotypes of either strain (data not shown); a result consistent with the notion that xylitol is internalized via a fructose-specific PTS (FruI) (18). In this case, our interpretation is that fructose competes better than xylitol for uptake, so less xylitol is internalized and there is less of an impact on growth.

**FIGURE 5.**
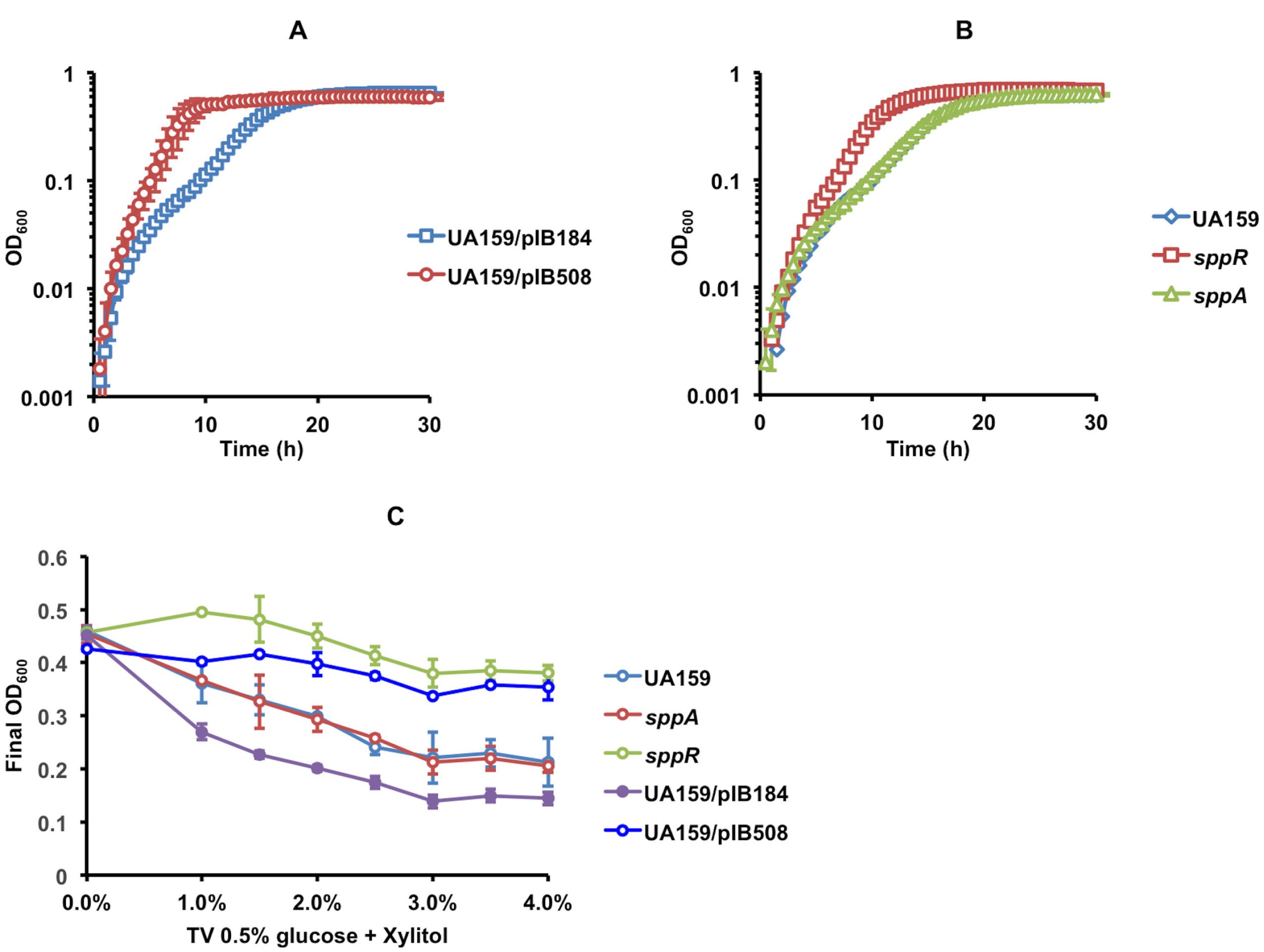
Overexpression of SppA enhances resistance to xylitol. Growth phenotypes of various strains were monitored over time in a Bioscreen C (A, B), or by measuring the final yield (OD_600_) after overnight incubation (C). Strains were cultivated first on BHI to exponential phase, then diluted using TV medium containing 0.5% glucose in addition to (A, B) 1% xylitol, or (C) the amounts of xylitol specified on the *x*-axis.

Since deletion of *sppR* enhanced the expression of *sppA* (Fig. 3B), we compared resistance to xylitol by the wild type and its otherwise-isogenic *sppR* and *sppA* mutants. The *sppR* mutant showed significantly improved growth in TV-glucose supplemented with xylitol, similar to UA159/pIB508 that overexpressed *sppA* (Fig. 5B&C). However, loss of *sppA* had no effect on xylitol resistance, as the *sppA* mutant grew similarly to the wild type. These results agree with our earlier findings that the expression of *sppA* is likely maintained at minimal levels, perhaps to avoid futile cycling. We have attempted, but have failed, to demonstrate the capacity of SppA to dephosphorylate xylitol-5-phosphate (X-5-P)(see Text S1 in the supplemental material), in large part due to the difficulty producing sufficient amounts of the compound; it is not commercially available.

### Accumulation of F-1-P enhances eDNA release and affects biofilm formation

*S. mutans* depends on the development of biofilms for its virulence, but little information is available on the effects of fructose on *S. mutans* biofilms. Wild-type cultures in exponential phase were grown in BHI and used to inoculate a biofilm medium (BM) constituted with 2 mM sucrose and 18 mM glucose (BMGS) or 18 mM fructose (BMFS), as well as a number of combinations of both glucose and fructose totaling 18 mM hexose. After 24 h or 48 h of incubation, the supernates of the culture were removed and development of biofilms was visualized and quantified by staining with 0.1% crystal violet. As shown in Fig. 6A, growth on fructose (BMFS) changed the accumulation of biomass by *S. mutans* on a polystyrene surface, resulting in a thinner but more uniform biofilm, as opposed to a patchy coverage by the biofilms formed on BMGS. Elution of crystal violet also indicated that greater amounts of dye were retained by the biofilms formed on fructose than glucose (Fig. 6A). This effect was not likely a result of reduced growth caused by a lack of glucose, as similar phenotypes were visible by the presence of as little as 3 mM fructose (data not shown). Also, to differentiate between effects of internalization of F-1-P versus F-6-P, we included the otherwise-isogenic mutants of UA159 deficient in either *fruI* (F-1-P) or *levD* (F-6-P). The results (Fig. 6A) showed that while the wild type and both mutants formed similar biofilms in BMGS, the *fruI* mutant formed patchy biofilms on BMFS, resembling those on BMGS. The *levD* mutant behaved like the wild type.

**FIGURE 6.**
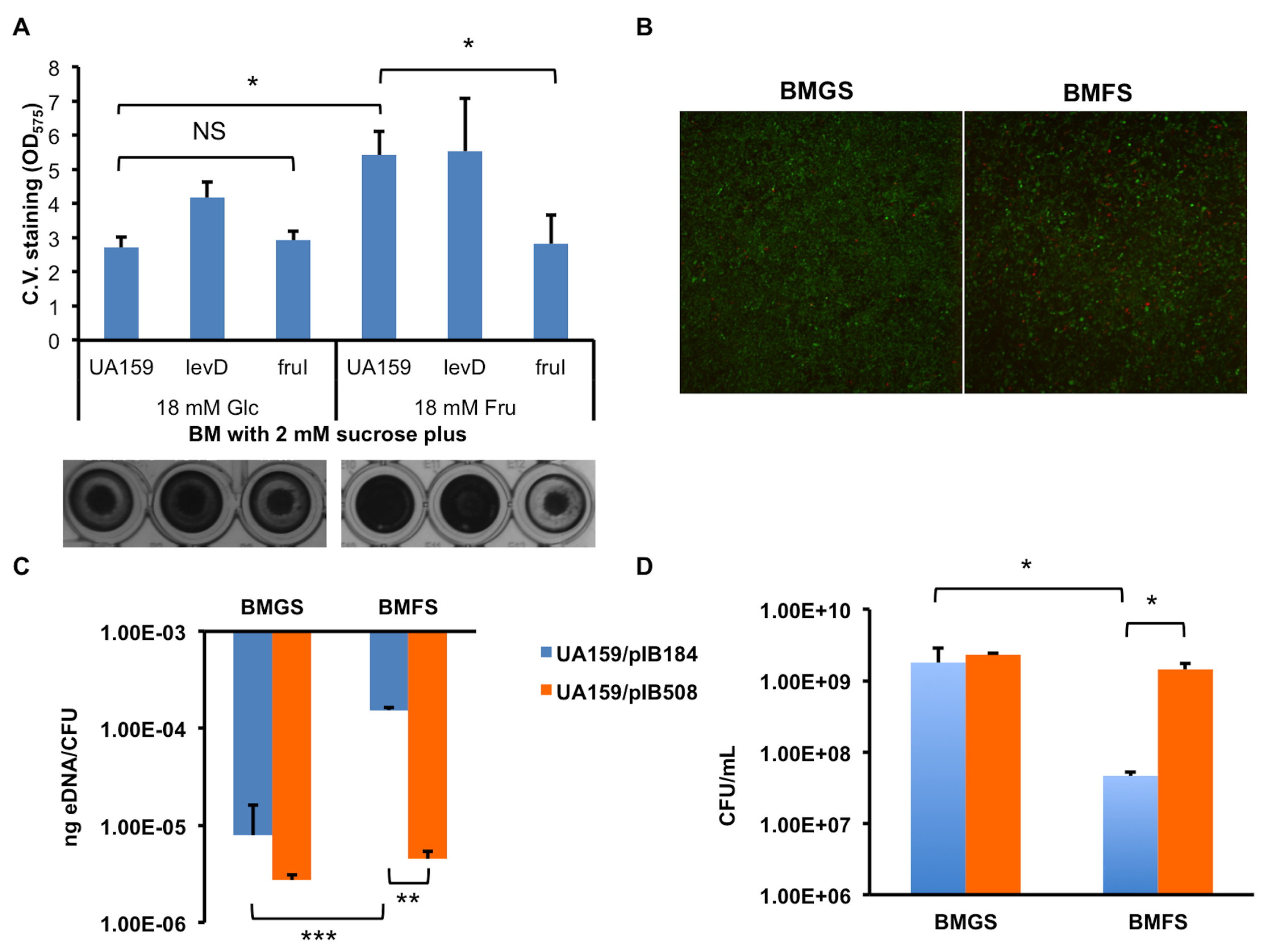
Fructose metabolism alters biofilm development by *S. mutans.* BM containing 2 mM sucrose and 18 mM glucose (BMGS) or 18 mM fructose (BMFS) was used for biofilm development. (A) Wild type UA159 and its isogenic mutants *levD* and *fruI* were each incubated for 24 h and the amounts of biofilms formed were visualized and quantified by crystal violet staining. (B) Biofilms formed by UA159 were treated with a LIVE/DEAD vitality stain and visualized under a confocal laser-scanning microscope, with green indicating live cells and red indicating dead cells. (C) eDNA released from 48-h biofilms formed by strains UA159/pIB184 (blue) and UA159/pIB508 (orange) were quantified using a DNA dye (SytoX Green) and a standard curve. The results were normalized using total CFU from each sample (D). Error bars represent standard deviations. Statistical analysis was performed using Student’s *t* test, with asterisks denoting *P* values: * <0.05, ** <0.005, and *** <0.0005.

Recent research has suggested that the ability of *S. mutans* to form efficient biofilms depends on both the synthesis of glucans and the release of extracellular DNA (eDNA) by bacteria, via both lysis-dependent and -independent pathways (30). To further study the effects of fructose, the release of eDNA was quantified using a fluorescent DNA dye in the biofilms of *S. mutans*. The results were normalized against the viable counts of bacteria (CFU). When grown as biofilms in BMFS, UA159 released significantly more eDNA per CFU after 24 h (5-fold) or 48 h (17-fold) of growth compared to BMGS medium (similar to UA159/pIB184 in Fig. 6C). Significantly lower CFU (5-fold for 24 h, and 20-fold for 48 h) were recovered from biofilms formed on BMFS than BMGS. As such, the amounts of eDNA detected per unit volume of supernatant fluid were comparable for both BMGS and BMFS. Therefore, it appears that the bacteria incubated in the presence of fructose experienced significantly more lysis, which resulted in increased release of eDNA and changes in formation of biofilms; although at this point we cannot rule out the involvement of other routes for production of eDNA. When treated with a LIVE/DEAD stain and observed under a confocal laser-scanning microscope, the biofilms formed on BMFS indeed showed notably more dead cells than those formed on BMGS (Fig. 6B). Similar impacts of fructose were observed for biofilm development by other genomically distinct, wild-type isolates of *S. mutans* (31), so the phenomenon is not restricted to the lab strain UA159 (Fig. S4 in the supplemental material). Furthermore, we compared the sensitivity of planktonic *S. mutans* cells to the cell wall-degrading enzymes lysozyme and mutanolysin by extracting chromosomal DNA after enzymatic treatment, using batch cultures grown in glucose- or fructose-containing medium. The results showed that 10-fold more DNA could be harvested from UA159 that was cultured in TV-fructose, compared to cells cultivated in TV-glucose, after each was treated with lysozyme, or a combination of lysozyme and mutanolysin, under otherwise-identical conditions (Fig. S5 in the supplemental material).

As a further test of our hypothesis that F-1-P in particular is responsible for the fructose effects observed so far, the strains UA159/pIB508 and UA159/pIB184 were compared in their ability to produce eDNA in the 48-h biofilm model. Similar to UA159, strain UA159/pIB184 yielded significantly lower numbers of CFU, but much higher levels of eDNA/CFU when growing on BMFS compared BMGS (Fig. 6C). When *sppA* was overexpressed on pIB508, significant reductions in the release of eDNA/CFU were apparent in comparison to strain UA159/pIB184, an impact that was especially significant for cells cultured in BMFS. At the same time, bacterial viable counts also showed significant recovery as an effect of pIB508, particularly when assayed using BMFS (Fig. 6D). These results suggest that accumulation of F-1-P in *S. mutans* likely leads to alterations in the cell-envelope that results in enhanced autolysis and release of eDNA, causing aberrant biofilm phenotypes.

### Other putative phosphatases capable of promoting sugar-phosphate resistance

The existence in *S. mutans* of a xylitol-inducible phosphatase capable of dephosphorylating X-5-P has been proposed (32). As xylitol reduced *spp* expression (Fig. 3A), we hypothesized that other phosphatase(s) are produced that may act on xylitol. Based on sequence similarity to SppA and other bacterial sugar-phosphate phosphatases, a functional genomics approach was employed to identify genes encoding products that could promote resistance to xylitol. A total of 9 ORFs (SMU.406c, SMU.428, SMU.488, SMU.718c, SMU.719c, SMU.742, SMU.1108c, SMU.1171c, and SMU.1830c) were selected and mutated by allelic exchange with an antibiotic resistance marker, and three (SMU.428, SMU.488 and SMU.718c) were chosen as potential candidates based on the decreased growth rates (Fig. S6 in the supplemental material) and yields (Fig. 7A) of the mutants in the presence of xylitol, compared to the wild type. For confirmation, these genes were expressed constitutively using pIB184, in both the wild-type and the respective mutant genetic backgrounds. When assayed for growth on TV-glucose supplemented with xylitol, mutants that were complemented by each of these three genes regained resistance to xylitol, and UA159 containing SMU.718c or SMU.488 expressed from pIB184 also showed enhanced yields, compared to their respective vector-only controls (Fig. S7 in the supplemental material). Furthermore, we previously reported that a deletion of the tagatose-1,6-bisphosphate aldolase (LacD) results in sensitivity to galactose in *S. mutans*, likely due to accumulation of metabolic intermediates that are normally catabolized by the tagatose pathway (33). When pIB428 was introduced into the *lacD* mutant, there was a significant alleviation of the inhibitory effects of galactose (Fig. 7B). We found that SMU.428, SMU.488 and SMU.718c are well conserved in the genomes of a diverse group of *S. mutans* isolates (31). Amino acid sequence alignments performed using these three proteins, together with SppA and the AphA phosphatase from *E. coli* (34), showed significant conservation in those amino acid residues (Fig. S8 in the supplemental material) that place these proteins in the family of bacterial class B acid phosphatases, part of the DDDD superfamily of phosphohydrolases (35). Computer analysis failed to identify any potential signal peptide sequences in the N-terminus of these proteins, suggesting they are not secreted. Further research is needed to understand their contribution to xylitol resistance and more broadly, to sugar-phosphate stress.

**FIGURE 7.**
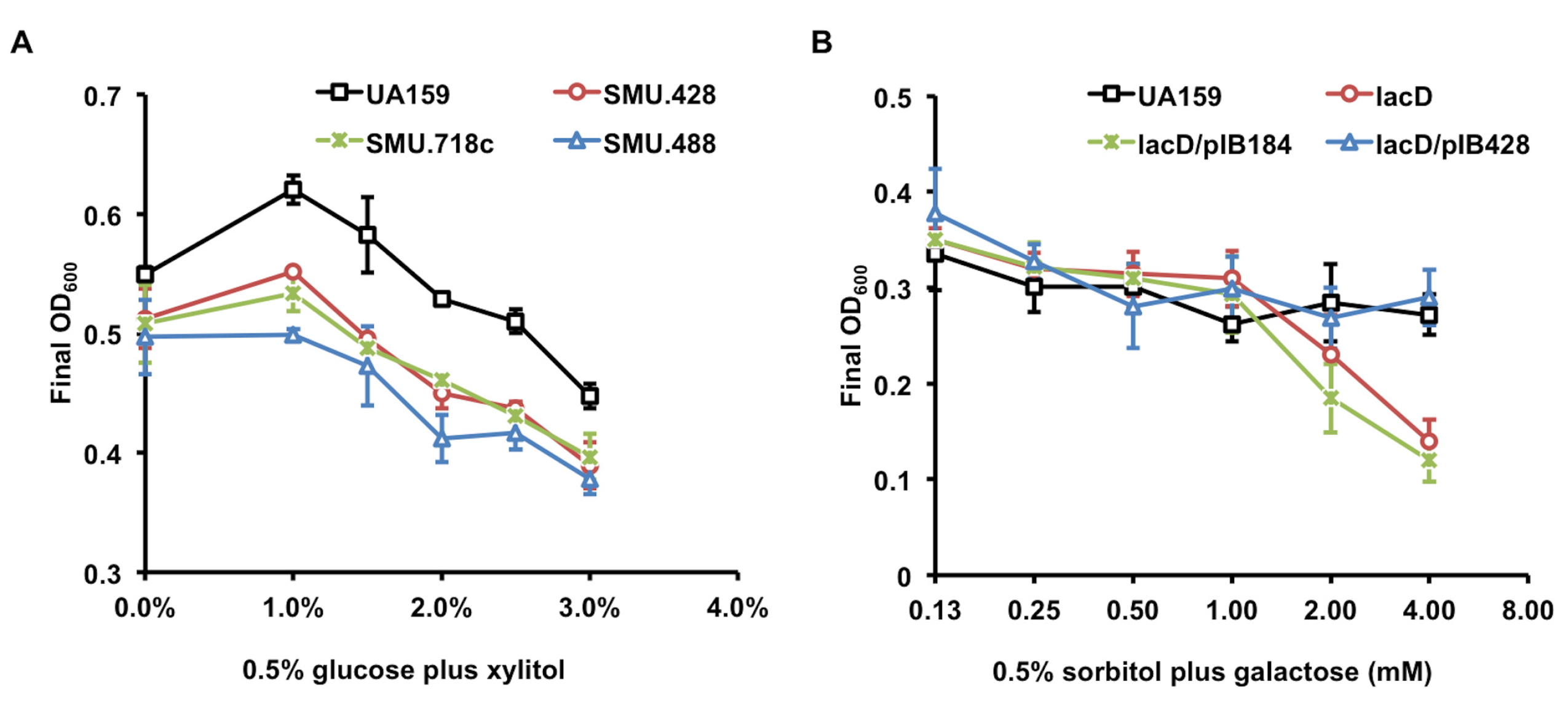
HAD family phosphatases contribute to sugar-phosphate tolerance. (A) *S. mutans* UA159 and its mutants deficient in putative phosphohydrolases. (B) UA159, a *lacD* mutant, and *lacD* derivatives containing plasmid pIB428 that overexpresses SMU.428 or the vector pIB184. Bacterial cultures were each prepared in BHI, then diluted into TV base medium supplemented with (A) 0.5% glucose and various amounts of xylitol or (B) 0.5% sorbitol and various amounts of galactose. After 17 to 20 h incubation at 37°C, OD_600_ of the cultures were recorded. Results are the averages of at least three independent cultures, with error bars denoting the standard deviations.

## Discussion

Carbohydrate fermentation is a highly dynamic process where the relative activities of various enzymes and resultant intermediates must be maintained at optimal levels to ensure fitness and survival, and to balance anabolic and catabolic processes. Perturbations to the systems, such as those that occur in the oral microbiome during intake of dietary carbohydrates by the host, require rapid responses by the microbiota. Sugar-related stress responses mediated by complex regulatory systems have been characterized mostly in Gram-negative model bacteria, e.g., *E. coli* (8, 36, 37). As a major caries pathogen, *S. mutans* depends on carbohydrate fermentation for energy generation and virulence. *S. mutans* is particularly well adapted to metabolizing sucrose, in part through the activities of three secreted glucosyltransferases (GtfB, GtfC and GtfD) that convert sucrose into glucan polysaccharides that serve as the scaffolding for biofilm accumulation and in creation of a diffusion-limiting matrix (38).

Interestingly, Gtfs generate a molecule of free fructose for every glucose moiety of sucrose that is targeted toward glucan formation. Furthermore, it is now clear that the majority of the sucrose presented to *S. mutans* is internalized by a sucrose-specific PTS permease (39). After internalization and phosphorylation, sucrose is cleaved by a sucrose-6-phosphate hydrolase into G-6-P and fructose. However, a significant portion of the fructose thus generated is expelled into the environment and re-internalized by fructose PTS permeases (26, 40). These observations, when coupled with the fact that fructose itself is a major constituent of foodstuffs, e.g. with high-fructose corn syrups increasingly used as common sweeteners, necessitate a better understanding of the metabolism of fructose by the oral microbiome.

*S. mutans* maintains at least three PTS permease operons (*fruRKI*, *fruPCD* and *levDEFG*) plus three phosphofructokinases (FruK, FruP, and SMU.1191/Pfk). Further, while a number of *S. mutans* PTS permeases can transport sugars other than their primary cognate carbohydrate, there are no other instances in *S. mutans* of redundancy in PTS permeases for sugars, besides those for fructose. This redundancy in PTS permeases, in itself, is noteworthy and could reflect significant evolutionary benefits to redundant fructose catabolic systems. Much remains to be clarified regarding the contributions of these PTS operons to the establishment and persistence of *S. mutans* in the human oral cavity, and to the organism’s cariogenic potential. However, it is understood that both F-1-P and F-6-P generated by the PTS are further phosphorylated to create F-1,6-bP, likely via 1-phosphofructokinase FruK and 6-phosphofructokinase SMU.1191, respectively. F-1,6-bP is then cleaved by an F-1,6-bP-specfic aldolase into glyceraldehyde 3-phosphate and dihydroxyacetone phosphate (3). We have demonstrated that removal of FruK leads to significant growth and fitness defects in *S. mutans*, apparently due to accumulation of the metabolic intermediate F-1-P (3). Here we present evidence that *S. mutans* maintains tight control over the levels of F-1-P through an inducible <u>s</u>ugar-<u>p</u>hosphate <u>p</u>hosphatase (*spp*) operon, consisting of a transcriptional regulator SppR and a fructose-phosphate hydrolase SppA. Changes in F-1-P levels, either increase or decrease, significantly alter bacterial physiology, including cell envelope integrity and biofilm development. We hypothesize that, aside from being an apparent stress compound in *S. mutans* (6), that F-1-P may also act as a signaling molecule, the accumulation of which promotes autolysis and release of extracellular DNA; with the latter being a critical contributor to formation of biofilms (30).

In support of this hypothesis, our latest RNA deep sequencing analysis in *S. mutans* has identified a fructose regulon that includes a significant number of genes that are likely involved in stress responses (26). For example, growth on fructose up-regulated the *cid* operon (SMU.1700c~1703c) that influences oxidative stress resistance and modulation of autolysis and biofilm development (41). Also up-regulated were the *rcrRPQ* and *relPRS* operons (SMU.921 to SMU.928) which have major impacts on acid and oxidative stress tolerance, competence development and (p)ppGpp metabolism (42). Thus, fructose and its intermediates are directly linked to pathways that are critical for persistence, growth and survival decisions, nutrient acquisition and genetic diversification (26).

On the other hand, a growing body of evidence suggests that F-1-P is acting as a regulatory molecule for *S. mutans*. Our recent study on the function of the *fruRKI* operon has detailed the transcriptomic impact of accumulation of F-1-P, a compound that likely serves as an allosteric effector to another DeoR-type regulator, FruR (3). It was clear that F-1-P influences gene regulation beyond the *fruRKI* operon. Specifically, the *fruK* mutant showed altered expression in as many as 20 known or probable transcriptional regulators, impacting ~20% of the genome. Key functions ranging from energy metabolism, protein and nucleic acid biosynthesis, to stress tolerance were affected by fructose. In addition to showing that overproduction of SppA, which preferentially degrades F-1-P, leads to phenotypic changes in both xylitol resistance and eDNA release, we have shown that the ability of fructose to induce *spp* operon expression depends on an intact FruI protein, which encodes the fructose-PTS permease that creates F-1-P (3). The ability of F-1-P to inhibit DNA-binding activity of the purified SppR protein *in vitro* provided direct evidence for its role as a regulatory effector. At the same time, the interaction of SppR with F-6-P that we observed also points to further complexity in this system. Notably, we have performed the same protein:DNA and protein:metabolite interaction assays using a different, His-tagged version of the SppR recombinant protein, and observed similar effects of both F-1-P and F-6-P on the binding of SppR to the promoter region of *spp* operon (data not shown). At first glance, F-6-P is unlikely the ideal signal molecule for fructose metabolism as it can also be generated from G-6-P through isomerization. However, F-6-P could act as a secondary allosteric effector for SppR under certain conditions, perhaps in a cooperative manner with F-1-P. Future research will focus on identifying other regulatory circuits that respond to the accumulation of F-1-P or F-6-P, both directly and indirectly, in particular the pathway(s) responsible for cell envelope integrity and biofilm phenotypes. As stated earlier, the *spp* operon has only been identified in a few oral streptococcal species, and its specificity for F-1-P could have implications in understanding niche adaptations and oral microbial ecology.

An intracellular hexose-6-phosphate:phosphohydrolase was purified from *Lactococcus lactis* that has a preference for galactose-6-phosphate and F-6-P (24). This enzyme had an estimated *M_r_* of 60 KDa by SDS-PAGE, was most active at pH 6.0, and its activity was stimulated by Mn^2+^, Mg^2+^ and Fe^2+^, and inhibited by fluoride. Another bacterial sugar phosphate phosphatase, Spp, was recently identified in *Pseudomonas fluorescens* that was capable of dephosphorylating ribose-5-P, F-6-P, and G-6-P (5). The optimal Mg^2+^ concentration for Spp was determined to be 2.5 mM, whereas addition of Mn^2+^ reduced the reaction rate. Therefore, SppA appears to share some similarities with other bacterial phosphohydrolases, yet it stands out with its preference for F-1-P. Lastly, despite our repeated efforts, we were not able to determine whether X-5-P is a substrate of SppA, partly due to our inability to obtain or generate sufficient X-5-P (it is not commercially available). As we continue to resolve this technical issue, it is important to note the possibility that SppA alleviates xylitol-related stress through its activities on either F-1-P or F-6-P, as X-5-P has been suggested to inhibit the activities of phosphofructokinases (9). The same rationale could apply in interpreting the xylitol phenotypes associated with the mutants that lacked one of three additional HAD phosphatases; that each enzyme may have multiple substrates and a stress phenotype could be alleviated by actions on molecules other than the one triggering the stress. Most notably, overexpression of SMU.428 abolished galactose-mediated stress in a *lacD* mutant. SMU.428 is located immediately downstream of the *copYAZ* operon that is responsible for copper efflux, biofilm formation and stress tolerance, yet its homologues in a number of other streptococci are closely associated with the tagatose operon. Clearly though, *S. mutans* possesses multiple phosphohydrolases and related regulatory proteins capable of mounting a stress response when triggered by certain sugar-phosphates. Additional research is needed to understand the function and regulation of these putative sugar-phosphate phosphatases.

## Materials and Methods

### Bacterial strains and culture conditions

*S. mutans* strains (Table 1) were maintained using brain heart infusion (BHI)(Difco Laboratories, Detroit, MI) agar plates or in liquid BHI medium at 37°C, in an ambient atmosphere supplemented with 5% CO_2_. To monitor bacterial growth phenotypes and for most other experiments, a semi-synthetic tryptone-vitamin (TV) medium (43) was utilized with various carbohydrates being provided at specified concentrations. Overnight bacterial cultures were prepared in BHI, then sub-cultured to exponential phase, and inoculated at 1:100 dilutions into TV media. Growth was monitored as the change in optical density at 600 nm (OD_600_) using a Bioscreen C lab system (Labsystems Oy, Helsinki, Finland). For development of biofilms in 96-well polystyrene plates or ibidi glass chambers (ibidi GmbH, Martinsried, Germany), a biofilm medium (BM)(44) was utilized that contained 2 mM sucrose and varying amounts of glucose or fructose, with bacterial inoculum prepared in BHI. Each of these experiments was repeated at least three times using independent cultures (biological replicates), with each sample being represented by at least two wells (technical replicates).

**Table 1.** Bacterial strains and plasmids used in this study.

| Strains or plasmids | Relevant genotypes<sup>a</sup> | Source or reference |
| --- | --- | --- |
| UA159 | Wild type, serotype c | (52) |
| FruK-13 | *fruKM28stop* | (3) |
| MMZ1387 | UA159 *sppR*::Km<sup>r</sup> | This study |
| MMZ1388 | UA159 *sppR*::Em<sup>r</sup> | This study |
| MMZ1399 | UA159 *sppA*::Km<sup>r</sup> | This study |
| MMZ1400 | UA159 *sppA*::Em<sup>r</sup> | This study |
| MMZ1416 | UA159 *P<sub>sppR</sub>*::*cat* | This study |
| MMZ1518 | *levD*::Em<sup>r</sup> *P<sub>sppR</sub>*::*cat* | This study |
| MMZ1519 | *fruI*::Sp<sup>r</sup> *P<sub>sppR</sub>*::*cat* | This study |
| MMZ1520 | *manL*::Em<sup>r</sup> *P<sub>sppR</sub>*::*cat* | This study |
| MMZ1521 | *ccpA*::Sp<sup>r</sup> *P<sub>sppR</sub>*::*cat* | This study |
| MMZ1546 | *sppR*::Km<sup>r</sup>/pBGE | This study |
| MMZ1547 | *sppR*::Km<sup>r</sup>/pBGE-*sppR* | This study |
| MMZ1549 | UA159 SMU.428::Km<sup>r</sup> | This study |
| MMZ1551 | UA159 SMU.718c::Km<sup>r</sup> | This study |
| MMZ1570 | UA159 SMU.488::Km<sup>r</sup> | This study |
| MMZ1573 | *sppR*::Em<sup>r</sup> *P<sub>sppR</sub>*::*cat* | This study |
| MMZ1574 | *sppA*::Em<sup>r</sup> *P<sub>sppR</sub>*::*cat* | This study |
| **Plasmids** |  |  |
| pIB184 | Shuttle expression vector, Em<sup>r</sup> | (46) |
| pBGE | Integration vector, Em<sup>r</sup> | (25) |
| pIB508 | *sppA* cloned into pIB184 | This study |
| pIB428 | SMU.428 cloned into pIB184 | This study |

<sup>a</sup> Km<sup>r</sup> kanamycin resistance; Sp<sup>r</sup>, spectinomycin resistance; Em<sup>r</sup>, erythromycin resistance.

**Table 2.**
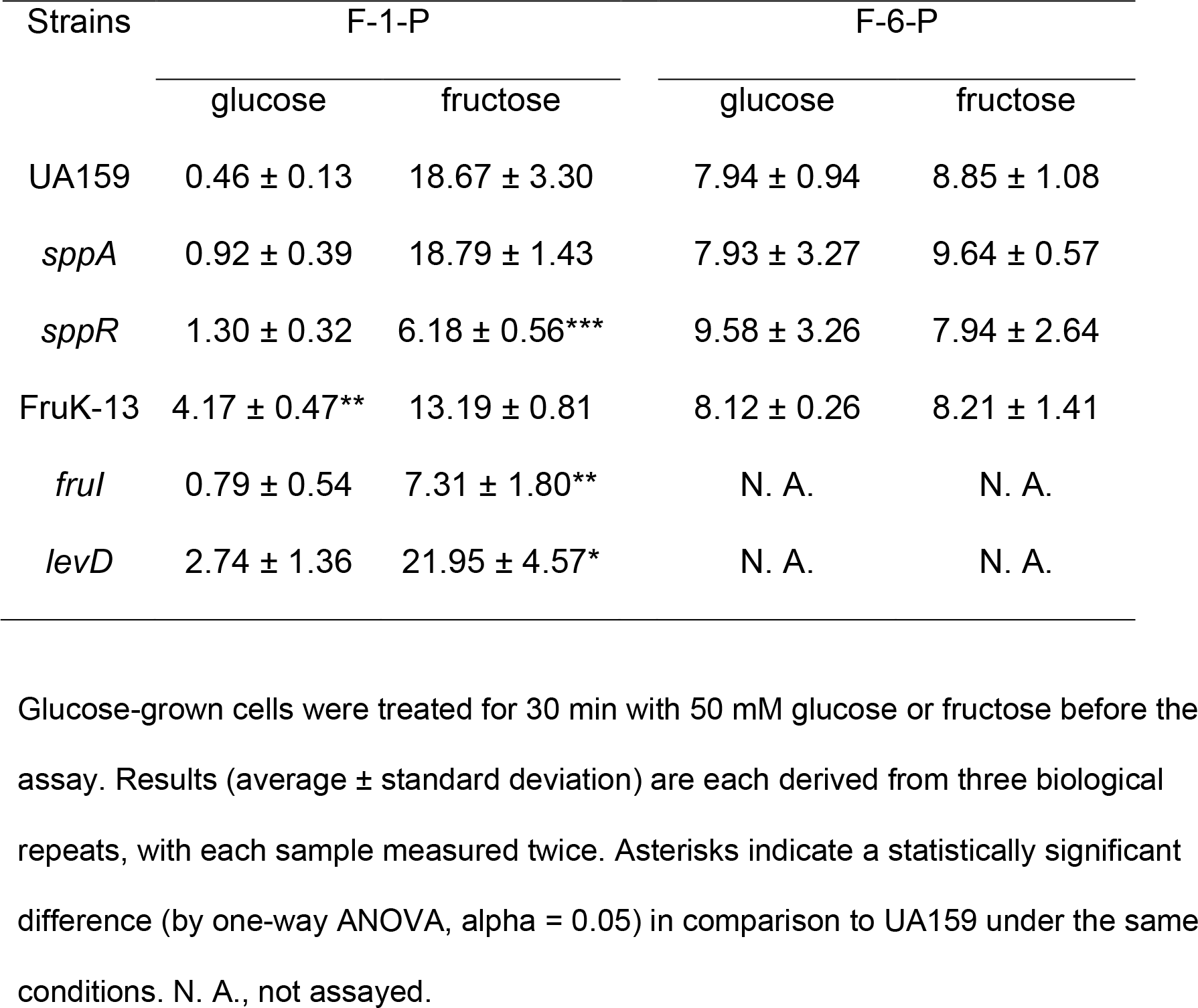
Measurements of fructose phosphates (nmole/mg protein).

| Strains | F-1-P |  | F-6-P |  |
| --- | --- | --- | --- | --- |
|  | glucose | fructose | glucose | fructose |
| UA159 | 0.46 ± 0.13 | 18.67 ± 3.30 | 7.94 ± 0.94 | 8.85 ± 1.08 |
| *sppA* | 0.92 ± 0.39 | 18.79 ± 1.43 | 7.93 ± 3.27 | 9.64 ± 0.57 |
| *sppR* | 1.30 ± 0.32 | 6.18 ± 0.56*** | 9.58 ± 3.26 | 7.94 ± 2.64 |
| FruK-13 | 4.17 ± 0.47** | 13.19 ± 0.81 | 8.12 ± 0.26 | 8.21 ± 1.41 |
| *frul* | 0.79 ± 0.54 | 7.31 ± 1.80** | N. A. | N. A. |
| *levD* | 2.74 ± 1.36 | 21.95 ± 4.57* | N. A. | N. A. |

Glucose-grown cells were treated for 30 min with 50 mM glucose or fructose before the assay. Results (average ± standard deviation) are each derived from three biological repeats, with each sample measured twice. Asterisks indicate a statistically significant difference (by one-way ANOVA, alpha = 0.05) in comparison to UA159 under the same conditions. N. A., not assayed.

### DNA methodology

Standard recombinant DNA techniques were employed in preparation of genetic materials for applications such as cloning and mutagenesis (45). To engineer a mutant defective in *sppR*, a set of 4 primers (see Table S1 for all primers used in this study) were designed for PCR amplification of two DNA fragments flanking the coding sequence of *sppR* (upstream fragment: SMU.507-1 and SMU.507-2GA; downstream fragment: SMU.507-3GA and SMU.507-4). PCR reactions were carried out using these primers to generate two DNA fragments that shared >25 bp overlaps with the 5’-and 3’- ends, respectively, of a kanamycin-resistance cassette. Subsequently, these flanking DNA fragments were used in equimolar concentrations, together with the kanamycin cassette, in a one-step ligation reaction that was catalyzed by a Gibson assembly kit (New England Biolabs, Beverly, MA). The ligated product was then used to transform a naturally competent UA159 culture, followed by selection on BHI agar plates containing kanamycin (1 mg/ml). Similar procedures were carried out in the creation of mutants defective in *sppA*, SMU.428, SMU.488, SMU.718c and other genes. All strains created in this study have been validated by DNA sequencing to ensure that no unintended mutations were introduced into the flanking regions of the gene of interest.

To construct a complementing plasmid for the *sppA* mutant, a pair of primers (SMU.508-5’Bm and SMU.508-3’SacI) were designed and used for amplification of the coding sequence of *sppA*, together with its putative ribosome-binding site. This DNA product was treated with restriction enzymes *Bam*HI and *Sac*I, and then ligated with an expression vector pIB184 (46) that was linearized with the same enzymes. After cloning into an *Escherichia coli* host strain BH10B, the resultant plasmid pIB508 was verified to contain the desired coding and regulatory sequences and introduced into the *sppA* mutant background, as well as the wild type UA159. Three additional plasmids, pIB428, pIB488, and pIB718c, were constructed in the same way, in the pIB184 vector, and used to complement their respective mutants. To complement the *sppR* mutant, the coding sequence of *sppR* was PCR-amplified (primers: sppR-5’XbaI and sppR-3’BsrGI) and cloned into an integration vector, pBGE, that places inserted DNA fragment under the control of the promoter of the *gtfA* gene (25). To monitor the transcriptional activities of the *spp* operon, a P*sppR::cat* fusion was constructed by cloning a DNA fragment containing the intergenic region upstream of *sppR* onto the integration vector pJL84 that contained a promoterless *cat* gene (47), and the gene fusion was subsequently delivered into a distal site on the chromosome of *S. mutans* by double cross-over recombination. The sequence information of all these primers is included in Table S1.

To assess the resistance of different UA159 cultures to muralytic enzymes, the bacteria were cultivated overnight in 5 ml TV medium containing 0.5% glucose or fructose, harvested by centrifugation at 4°C and washed once with a TE buffer (10 mM Tris-Cl, 1 mM EDTA, pH 8.0). Cells were resuspended in 500 μl of lysis buffer that contained 50 mM Tris-Cl at pH 8.0, 10 mM EDTA, 10 mg/ml of lysozyme, and 0.1 mg/ml of RNase A. After addition of 0 or 20 units of mutanolysin (Millipore Sigma, St. Louis, MO), cells were incubated at 37°C for 1 h, and lysed by adding 40 μl of 20% SDS and incubating at 80°C for 15 min. Subsequently, 80 μl of 5 M NaCl was added into each reaction, mixed well, followed by addition of 600 μl of phenol: chloroform: isoamyl alcohol (25: 24: 1, v/v/v) and 2 min of vortexing. The solution was then clarified by centrifugation at room temperature at 15000 × *g* for 5 min, followed by the removal of 500 μl of the aqueous phase. The genomic DNA was then precipitated from the aqueous solution by addition of 500 μl 100% isopropanol and centrifugation as before. After washing once with 75% ethanol, the DNA was air-dried and dissolved in 50 μl TE buffer and quantified using a Nanodrop apparatus (Thermo Fisher, Waltham, MA) and agarose-gel electrophoresis.

### Engineering and preparation of recombinant proteins of SppR and SppA

DNA fragments containing the entire coding sequences of SppR and SppA were each amplified using PCR primers (SMU.507-5’Bm and SMU.507-3’HIII for *sppR*, and SMU.508-5’Bm and SMU.508-3’HIII for *sppA*)(Table S1) containing engineered restriction sites, and digested using restriction enzymes *Bam*HI and *Hin*dIII. After ligating with a pQE30 vector prepared by treating with the same enzymes to create an N-terminal, 6× His-tagged SppA, the construct was introduced into *E. coli* M15. Similarly, the digested product of *sppR* was cloned into a plasmid vector pMAL-p2X and expressed as a fusion protein (MBP-SppR) in *E. coli* NEB Express. In order to produce a recombinant 1-phosphofructokinase for the F-1-P assay (below), an *E. coli* Fru1PK gene was also cloned into pQE30, using primers 1-PFKEcoli-5Bm and 1-PFKEcoli-3PstI (48). After confirming the fidelity of the PCR and cloning via DNA sequencing, the expression of each recombinant protein was induced by adding IPTG to a final concentration of 0.05 mM to cells growing at mid-exponential phase.

Purification of recombinant proteins, MBP-SppR and His-SppA, was carried out using an amylose resin (New England Biolabs) and an Ni-NTA agarose (Qiagen), respectively, according to protocols provided by the suppliers, with some modifications. Briefly, to improve the stability of MBP-SppR, 10% of glycerol and 0.3 M of NaCl were supplemented to the original column buffer (20 mM Tris-Cl, 0.2 M NaCl, and 1 mM EDTA, pH 7.5). After elution from the column, MBP-SppR was treated with Factor Xa (New England Biolabs) to release the SppR protein, and the products were dialyzed three times, at a 1:1000 dilution, against a buffer containing 10 mM HEPES pH 7.5, 0.5 M NaCl and 20% glycerol, using a Slide-A-Lyzer dialysis device with a MWCO of 3.5 KDa (Thermo Fisher). To avoid carryover of phosphate into the subsequent reactions involving the recombinant SppA, purification of His-SppA was performed using a phosphate-free buffer system that was created by replacing NaH_2_PO_4_ in the original buffers with 50 mM HEPES (pH 7.9), and also by supplementing with 10% glycerol. Dialysis of the purified His-SppA solution was carried out against 10 mM PIPES pH 6.5, 0.5 M NaCl, and 20% glycerol.

### Electrophoretic mobility shift assay (EMSA) and thermal shift

To study the interaction of SppR with the promoter of *sppR*, EMSA was performed using a biotin-labeled DNA fragment containing the intergenic region upstream of the *sppR* coding sequence as previously described (49). When needed, various sugar phosphates were each used at 5 or 10 mM. To study the interaction of SppR with various sugar phosphates, a 10-μM SppR solution was prepared using a buffer containing 0.1 M HEPES (pH 7.5) and 0.15 M NaCl. After addition of a SYPRO Orange protein stain (Thermo Fisher), used at a 300-fold dilution, and various sugar phosphate at specified amounts, the 10-μL reactions were subjected to a melting-curve treatment on a CFX96 Touch real-time PCR thermocycler (Bio-Rad, Hercules, CA) by increasing the temperature from 25°C to 75°C in increments of 0.2°C. The RFU measurements were then plotted against the temperature values to determine the melting temperature (Tm) (29).

### Phosphatase assay

The ability of His-SppA to remove the phosphate group from various sugar-phosphates was measured at 37°C. The basic buffer was composed of 50 mM PIPES pH 6.5, 5 mM MgCl_2_, and 1 mM MnCl_2_. For other pH values, different buffering reagents were used in place of PIPES at the same concentration: 2-ethanesulfonic acid (MES) or potassium citrate (K-Cit) for pH 5.5; 1,4-piperazinediethanesulfonic acid (PIPES) for pH 6.0 and pH 6.5; and HEPES for pH 7.0 and pH 7.5. All initial experiments to determine the effects of metal ions were carried out in an end-point assay, using HEPES-based buffers prepared at neutral pH.

To measure the reaction kinetics of His-SppA on F-1-P, 20 μL of His-SppA that was diluted to 1 μM using the dialysis buffer was mixed with 40 μL of F-1-P (Santa Cruz Biotechnology, Dallas, TX) prepared in the basic buffer, at final concentrations ranging from 0.1 mM to 1.0 mM; for F-1-P ranging from 1.5 mM to 10 mM, 0.25 μM of protein was used. All components were pre-warmed to 37°C before mixing. With the reaction maintained at 37°C, a 5-μL aliquot of the reaction was removed at 1 min intervals and subjected to measurement of free phosphate using a phosphate assay kit (Malachite Green Reagent, MGR; Sigma). For the kinetics of His-SppA with F-6-P as a substrate, 20 μL of His-SppA at 2 μM was used in each reaction, and aliquots of the reaction were taken and measured every 5 min. Each reaction was repeated twice. The data were analyzed using the Prism from GraphPad software (La Jolla, CA).

### F-1-P and F-6-P assays

Bacteria were cultivated overnight in TV with 10 mM glucose, then diluted 1:20 ratio into 50 mL of the same medium and sub-cultured until the OD_600_ reached 0.3 ~ 0.4. 50 mM glucose or fructose was then added into the culture. After another 30 min of incubation, cells were harvested by centrifugation at 4°C, immediately resuspended in 600 μL PBS, and mixed with 500 μL of glass beads. After two rounds of homogenization at 4°C, cell lysates were clarified by centrifugation (16,000 × *g*, 10 min, 4°C). A BCA assay (Thermo Fisher) was carried out to quantify the protein concentrations. Deproteinized lysates were generated by mixing 400 μL of cell lysate with 100 μL of 100% (w/v) trichloroacetic acid and incubating on ice for 10 min. After centrifugation, 420 μL of the supernate was removed and neutralized by adding 50 μL of 6 N NaOH, and the pH value of each sample was confirmed using a pH-indicator strip. The deproteinized cell lysate was used immediately for enzymatic assays or stored at −80°C.

Each 1 mL master mix of the F-1-P assay included 20 mM HEPES, pH 7.1, 50 mM KCl, 5 mM MgCl_2_, 1 mM DTT, 2.5 mM ATP, 0.05 mM NADH, 1 μL glycerol-3-P dehydrogenase (Sigma), 10 μg triose-P isomerase (Sigma), 10 μg of rabbit muscle aldolase in ammonium sulfate suspension (Sigma), and 5 μL of His-1PFK (18.6 μg/μl) that was engineered based on the *E. coli* 1-phosphofructokinase gene. 10 μL of deproteinized cell lysate was mixed with 90 μL of the master mix. After 15 min of incubation at 37°C, optical density OD_340_ was measured using a spectrophotometer DU640 and a quartz cuvette, and the F-1-P levels were determined against a standard curve prepared using F-1-P at concentrations ranging from 1 to 20 μM. A control reaction was included for each sample by omitting the His-1PFK protein, and the measurements were subtracted from the actual results. Each strain was represented by 3 biological replicates, and each sample was measured at least twice. Levels of F-6-P in the same lysates were measured using a F-6-P assay kit (Sigma) by following protocols provided by the supplier.

### Biofilm development, microscopy and measurement of extracellular DNA (eDNA)

Development of biofilms by *S. mutans* was investigated in 96-well polystyrene plates (Thermo Fisher) for comparison of overall biofilm mass, or on glass surfaces using an ibidi 8-well μ-Slide to assess structural changes and release of eDNA. A semi-defined biofilm medium (BM) was modified to contain final concentrations of 2 mM sucrose and 18 mM glucose (BMGS) or fructose (BMFS)(3, 50). Exponential-phase bacterial cultures prepared using BHI were diluted 1:100 into BMGS or BMFS and incubated at 37°C in an aerobic environment supplemented with 5% CO_2_. Each experiment included at least three biological replicates. The 96-well plates were incubated for 24 to 48 h, washed with phosphate-buffered saline (PBS), and stained with 0.1% crystal violet (CV). After extensive washing with water, the retained CV was eluted with a solution of 30% acetic acid and quantified by measuring the optical density at 575 nm (OD_575_).

The ibidi slides were incubated for 48 h, with a replacement of medium after first 24 h. For observation of biofilm structures, each biofilm was treated using a LIVE/DEAD BacLight bacterial vitality stain kit (Thermo Fisher) and visualized using a confocal laser scanning microscopy system (4). To assess the amount of eDNA released by each biofilm, the entirety of the 48-h biofilm culture was harvested by scraping off the biofilm material, without washing, and sonicated for 30 sec at 100% power using a water-bath sonicator (model FB120, Thermo Fisher). A portion of the cell suspension was subjected to serial dilutions and plated on BHI plates for enumeration of CFU, and the remaining material was pelleted using a tabletop centrifuge for 2 min at 16,000 × *g*. Subsequently, a portion of the supernatant fluid was removed and combined 1:1 with a 5 μM solution of SYTOX Green nucleic acid stain (Thermo Fisher), followed by measurement of fluorescence (excitation/emission: 504/523 nm) using a fluorescence-capable Synergy 2 plate reader from BioTek (Winooski, VT). A standard curve was established using a known DNA solution and the results were calculated as ng of eDNA released per CFU of the biofilm.

### Quantitative reverse transcription PCR (qRT-PCR)

Total RNA was extracted from *S. mutans* cells harvested at OD_600_ ≈ 0.5 using an RNeasy kit (Qiagen) by following an established protocol (51). Primers specific to *sppA* were used for both the reverse transcription and Real-time PCR that followed, with *gyrA* serving as an internal control (26).

### CAT (*c*hloramphenicol *a*cetyl*t*ransferase) assay

To assess the expression of P*sppR::cat* fusion in various backgrounds, specific CAT activities from clarified bacterial lysates were measured according to an established protocol detailed elsewhere (47).

## Acknowledgment

This work was supported by DE12236 from the National Institute of Dental and Craniofacial Research.

